# Structural Adaptations of HepI Enzymes in Proteobacteria: Insights into Evolutionary Resilience and Functional Dynamics

**DOI:** 10.1101/2024.09.01.610683

**Authors:** Aayatti Mallick Gupta, June W. Butelman, Bakar A. Hassan, Philip Arevalo, Erika Anne Taylor

## Abstract

Since lipopolysaccharide molecules are highly variable in their structure while also being essential for bacterial cell survival, we hypothesized that enzymes involved in its biosynthesis may exhibit differences that could be used to predict the likelihood of survival in different ecological niches. We examined the sequence variability of orthologues of Heptosyltransferase I (HepI), which is found in all lipopolysaccharide (LPS) containing Gram-negative bacteria. We identified two different sequence motifs within the N-terminal domain of HepI, which correlated with differences in the Lipid A portion of the LPS. Further, we compared the protein structure of HepI from *Escherichia coli* with a structural model we generated that incorporated the alternate sequence motif. Molecular dynamics simulations of these two proteins, recapitulated our findings, that proteins with the *E. coli*-like sequence motif maintained a larger enzyme active site, while the mutated structural model undergoes rearrangements that lead to a smaller N-terminal active site pocket. This work revealed sequence-structure-function relationships that can be used to determine if a species incorporates a Heptose residue onto Lipid A chains containing one or two 3-deoxy-D-*manno*-oct-2-ulosonic acid (Kdo) residues. Since research suggests that the number of Kdo residues in LPS impacts the overall immune response to the LPS endotoxin, this work could aid in our understanding of the pathogenic effects of human-bacterial interactions. Understanding the sequence-structural adaptations of HepI enzymes across proteobacteria sheds light on their evolutionary resilience and functional versatility.

**Highlights:**

- Lipopolysaccharide (LPS) structure varies between organisms
- Heptosytransferase I sequence motifs allow prediction of LPS structural features
- Adaptations of HepI enzymes reveal evolutionary resilience and functional versatility

## Introduction

(The introduction should clearly state the objectives of your work. We recommend that you provide an adequate background to your work but avoid writing a detailed literature overview or summary of your results.)

Lipopolysaccharides (LPS), integral components of the outer membrane in Gram-negative bacteria, play pivotal roles in environmental adaptation and pathogenicity [1]. The biosynthesis of LPS exhibits considerable diversity among proteobacteria, influenced by environmental pressures that drive subtle yet significant modifications in their structural and functional properties [2]. Central to this biosynthetic pathway is Heptosyltransferase I (HepI), a key enzyme responsible for building the core component of LPS which is crucial for bacterial membrane integrity and host interaction. HepI catalyzes the incorporation of the L-*glycero*-D-*manno*-heptose (Hep) moiety onto the first Kdo (Kdo: 3-deoxy-D-*manno*-oct-2-ulosonic acid) of Lipid A in Gram-negative bacteria [3–7] (Figure 1a). HepI enzymes across proteobacteria exhibit sequence motifs that respond to structural variations in lipid A, including attributes such as hexosamine composition, phosphorylation levels, and acyl chain characteristics. This sequence heterogeneity underscores the adaptability of proteobacteria to diverse environmental niches, where specific sequence motifs reflect evolutionary strategies tailored to ecological challenges. To understand the role of HepI variability in conferring the necessary diversity in cell membrane properties to allow adaptation to different ecological niches we generated multiple sequence alignments (MSAs) to identify patterns in the sequence-structure-function variability.

**Figure 1:**
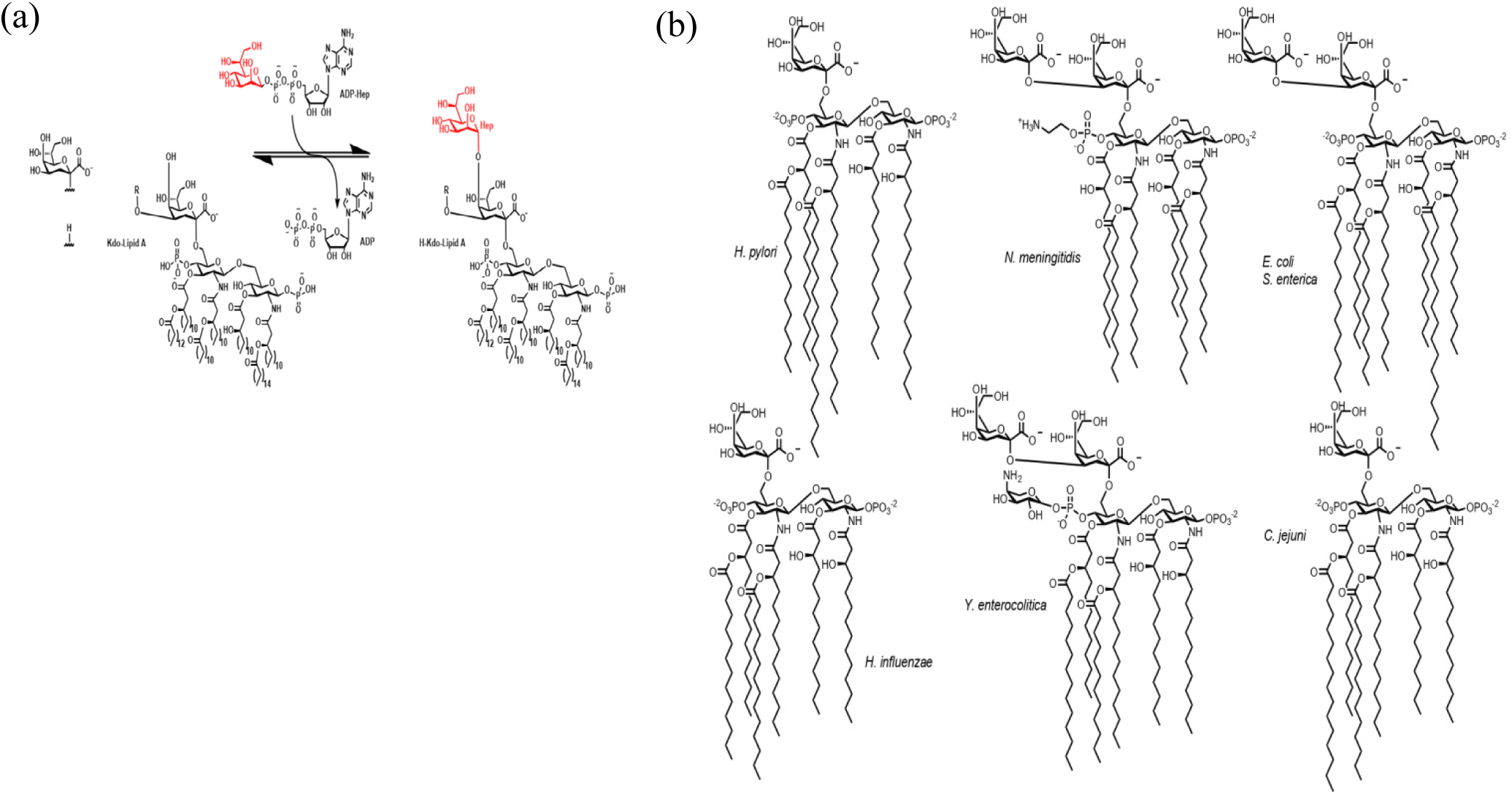
(a) The transference reaction typically catalyzed by wt-HepI. L-glycero-D-manno-heptose is transferred from a donor substrate, ADP-heptose, to acceptor substrate H-Kdo-Lipid-A. (b) Lipopolysaccharide (LPS) variants differ across various proteobacteria, reflecting their diverse structural and functional adaptations.

It has been observed γ proteobacteria exhibit a uniquely flexible nature in their evolutionary trajectory, which allows them to adapt to a wide range of environmental stresses, showcasing shared adaptive responses that are observed across various proteobacterial classes [8]. Within γ proteobacteria, HepI sequences exhibit distinct clades such as Enterobacteriales and Pseudomonales display diverse sequence motifs that blend characteristics of both α/β and ε/δ/ζ proteobacteria. This indicates convergent evolution or functional conservation shaped by their ecological roles, further underscoring their adaptability and versatility. Further examination of the HepI MSA revealed differential sequence conservation patterns in residues known to be located within the active site of the N-terminal domain Lipid A binding domain. Examination of these motifs in combination with analyses of LPS variants (Figure 1b), revealed that bacterial species that incorporate only one Kdo residue on Lipid A, such as *Haemophilus influenza*, maintained one sequence motif, while LPS from species that incorporate two Kdo residues on Lipid A, such as *Escherichia coli* had a different conserved sequence motif. Due to the lack of a HepI protein crystal structure for *H. influenza*, we generated a *Haemophilus*-like version of the *E. coli* protein structure (mt-HepI) which we used for comparison to the *E. coli* (wt-HepI) structure (PDB ID: 2GT1). This comparison revealed intriguing insights into how changes in the enzyme’s active site affected substrate binding and enzymatic selectivity. We hypothesized that *Haemophilus*-like mt-HepI enzymes would exhibit a smaller binding pocket tailored to accommodate the single Kdo containing Lipid A substrate, in contrast with those *Escherichia*-like wt-HepI enzymes which would have a larger, more flexible binding pockets to accommodate Lipid A molecules with two Kdo sugars.

These two structural variants were used for 1 microsecond molecular dynamic simulations to enable examination properties such as radius of gyration (Rg), solvent-accessible surface area (SASA), and dynamic cross-correlation matrices (DCCM), to provide a comprehensive view of how these different sequence motifs reshaped HepI’s substrate specificity. Reductions in Rg and SASA, coupled with increased β sheet formations and altered residue interactions, highlighted adaptive responses that stabilized mt-HepI proteins, ensuring enzymatic functionality despite structural perturbations to select for utilization of a smaller substrate.

## 2. Materials and Methods

### 2.1 Multiple sequence analysis

Heptosyltransferase I (HepI) sequences (Table S1) from various classes of Gram-negative proteobacteria was retrieved from InterPro [9]. Multiple sequence alignments (MSAs) of HepI from these different classes were generated using MAFFT with the FFT-NS-i algorithm [10] and Clustal Omega [11]. These alignments illustrated the sequence variability of HepI across Gram-negative proteobacteria. Specifically, this study emphasized the N-terminal sugar acceptor substrate binding region to provide a detailed comparison of sequence diversity among these bacterial classes.

### 2.2 Selection of Sequences for Phylogenetic Analysis

A set of HepI sequences from various species of proteobacteria was obtained by searching for matches to KEGG proteins K19282 (heptosyltransferase I) and K02841(lipopolysaccharide heptosyltransferase I) in Annotree (Table S1) [12]. These sequences were aligned using MAFFT with the FFT-NS-i algorithm [10] and used to construct a maximum likelihood phylogenetic tree with IQ-TREE. HepII sequences from were used as an outgroup to root the tree [13]. This tree was visualized using iTOL [14] and used to investigate the evolution of HepI by comparing it to the phylogenetic tree of reference species generated by the Genome Taxonomy Database (Figure S1) [15].

### 2.3 Modeling

The X-ray crystallographic structure of *E. col i* HepI (2GT1.pdb), referred to as wt-HepI, served as the reference model. To study the structural impact of specific mutations, a structural model of *H. influenzae*-like HepI (mt-HepI) was created. In this model, the S10A mutation was introduced at position 10, altering the sequence to SAXGD from positions 9 to 13. This modification was followed by incorporating the conserved region VVYDK at positions 60 to 64.

The structural modifications were performed using Discovery Studio Visualizer. This software allowed for precise manipulation of the protein structure, enabling the introduction of the S10A mutation and the adjustment of the sequence to reflect the SAXGD and VVYDK motif, ensuring that the structural model accurately represented the sequence-structure relationship of *H. influenzae* HepI. This approach allowed for detailed comparative analysis with the wt-HepI structure from *E. coli*, providing insights into the structural adaptations and potential functional implications of the HepI variants.

### 2.4 Simulation System Setup

Comparative molecular dynamics (MD) simulations were conducted to explore the structural impact of HepI variants across different proteobacterial groups, specifically representing the structures of *E. coli* and *H. influenzae*. We conducted all-atom MD simulations under standard isothermal-isobaric ensemble (NPT) conditions at 310 K and 1 atm pressure, using the GROMACS 2018.6 package [16], with periodic boundary conditions, the spc216 water model, and the GROMOS 53a6 force field [17]. The protein was solvated in a simulation box with Simple Point Charge (SPC) water molecules [18] and neutralized by adding six chlorine ions. Energy minimization was performed using the steepest descent method with a 100 kJ/mol tolerance. Van der Waals and electrostatic interactions were identified with cutoffs of 1.4 nm and 1.2 nm, respectively, using the Particle Mesh Ewald (PME) method [19]. The LINCS algorithm constrained all bond lengths, and the SETTLE algorithm constrained water molecule geometry. The equilibration process included a 100 ps NVT run at 300 K with a 0.1 ps coupling constant, followed by a 100 ps NPT run at 1 bar with a 5 ps coupling constant, utilizing the Berendsen coupling scheme. The final 1 μs production run was conducted for both wildtype and mutant HepI to identify structural changes in the N-terminal substrate acceptor site, maintaining consistent conditions across all systems to ensure equivalent simulated ensembles.

### 2.5 Trajectory analysis

The structural behavior of wt-HepI and mt-HepI proteins in *E. coli* and *H. influenzae* was analyzed using molecular dynamics (MD) simulations. Trajectory files were used to compare the structural dynamics of these proteins. Analogous to prior HepI MD simulations, critical structural analyses, including RMSD, RMSF, Rg, hydrogen bond and SASA, were performed using the standard protocol of GROMACS package [20–22]. Intermolecular hydrogen bonds (protein-water interactions) were calculated with NH bond default values set to less than 0.35 nm for donor-acceptor distances. For each frame of the selected trajectories, we calculated the center of mass for the residues within regions 10-13, 60-63, and 120. We then measured the Euclidean distances between these centers of mass and recorded the minimum distance for each frame to create a time-resolved profile of the spatial relationships between the active sites. The distance measurements were averaged over the selected frames to obtain representative values for both the wildtype and mutant proteins, allowing for a comparative analysis of the spatial arrangement of the active sites. DSSP calculations have been performed using the GROMACS interface which allows for detailed analysis of secondary structure changes during a molecular dynamics simulation. This integration helps in understanding the effects of mutations, ligand binding, or environmental changes on protein structure and stability. All the analysis was performed using the conserved trajectory (Figure S2) from 700 ns till the end of the simulation.

### 2.6 Binding Pocket Area and Volume Analysis

We utilized CASTp (Computed Atlas of Surface Topography of proteins) to examine the binding pocket area and volume [23]. CASTp is an advanced tool that identifies and measures surface pockets and interior voids within protein structures. To capture the steady-state conformations of both the wildtype and mutant HepI proteins, we selected structures from the converged portion of the simulation, particularly between 700 and 1000 ns. By sampling structures at 50 ns intervals, we obtained a representative set of 7 structures for each protein variant. The selected structures underwent preparation for CASTp analysis. This involved removing all water molecules and ions, ensuring that only the protein coordinates were included in the final PDB files. This step was crucial for accurate binding pocket analysis.

### 2.7 Principal Component Analysis

The advanced analysis of the trajectories from MD simulations was performed using Essential Dynamics (ED) or Principal Component Analysis (PCA). Through PCA, the concerted motion of the protein was identified by analyzing different frames during the simulations. The PCA process involved two main steps: (1) constructing a variance/covariance matrix using C-α atoms and (2) diagonalizing the covariance matrix. The covariance matrix was built from atomic fluctuations after removing translational and rotational movements, allowing the identification of the principal 3N directions along which most protein motion occurs. This matrix was represented as a simple linear transformation in Cartesian coordinate space. The diagonalization of the data covariance (or correlation) matrix C in PCA was achieved by computing C=VKV^T^, where K contains the eigenvalues as diagonal entries and V contains the corresponding eigenvectors. The values of C range from −1 to 1. A positive value represents positively correlated movement between the iiith and jjjth residues, while a negative value indicates negatively correlated movement. PCA was implemented on the position covariance matrix, constructed using atomic coordinates from the MD trajectory. To investigate the dynamic behavior and structural correlations of heptosyltransferase I (HepI) from *E. coli* (wildtype) and *Haemophilus influenzae* (mutated), we utilized PCA and Dynamic Cross-Correlation Matrix (DCCM) calculations, facilitated by the Bio3D package in R.

## 3 Results

### 3.1 Conserved Substrate Binding Motifs in HepI: Sequence and Phylogenetic Analysis

We examined the N-terminal substrate acceptor site of the GT-B fold in HepI from different Gram-negative proteobacteria, since this is the region that recognizes differences in Lipid A structure [24]. Multiple sequence alignment (MSA) of HepI proteins exported from InterPro revealed noticeable differences in the amino acid composition of residues previously identified to be important for ligand interactions [21, 25, 26] including those corresponding to *E. coli* HepI residues 9-13 and 60-64 (data not shown). An MSA was then generated from a group of representative taxonomic classes of proteobacteria indicated characteristic amino acid sequence features in the conserved substrate binding site (Figure 2). α and β proteobacteria exhibited the distinctive SSXGD conserved substrate binding motif at amino acid positions 9-13, along with the RRWRK conserved region at positions 60-64. ε, δ, and ζ proteobacteria typically showed a S10A mutation, resulting in the SAXGD motif, followed by the VVYDK/IVYDK conserved region at positions 60-64. γ proteobacteria displayed dual features: Enterobacteriales and Pseudomonales exhibited α and β proteobacteria-like conserved motifs in the 9-13 and 60-64 substrate binding regions, while the rest of the γ proteobacteria elicited motifs similar to ε, δ, and ζ proteobacteria.

**Figure 2:**
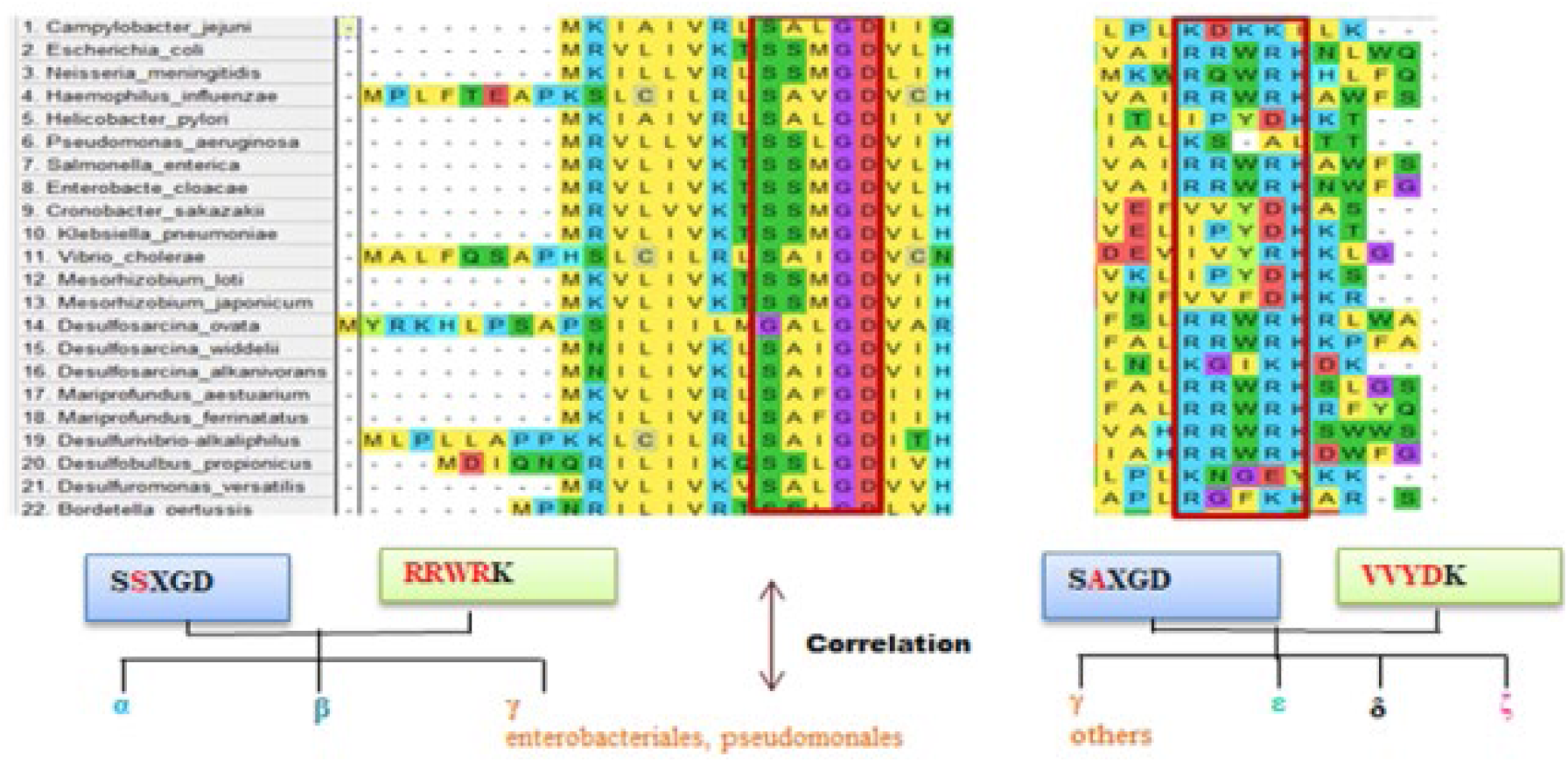
Multiple sequence alignment (MSA) of HepI from representative taxonomic classes of proteobacteria.

Detailed phylogenetic analysis of HepI sequences revealed distinct evolutionary clades among various classes of proteobacteria (Figure 3). Specifically, HepI sequences from Enterobacteriales and Pseudomonales, subgroups within γ proteobacteria, formed distinct clades that separated from the rest of the proteobacteria. While Enterobacteriales and Pseudomonales shared some sequence motifs with α and β proteobacteria, they also showed features akin to ε, δ, and ζ proteobacteria. This dual characteristic suggested that γ proteobacteria possess a unique evolutionary versatility, allowing them to adapt and evolve distinct sequence features in response to diverse environmental pressures.

**Figure 3:**
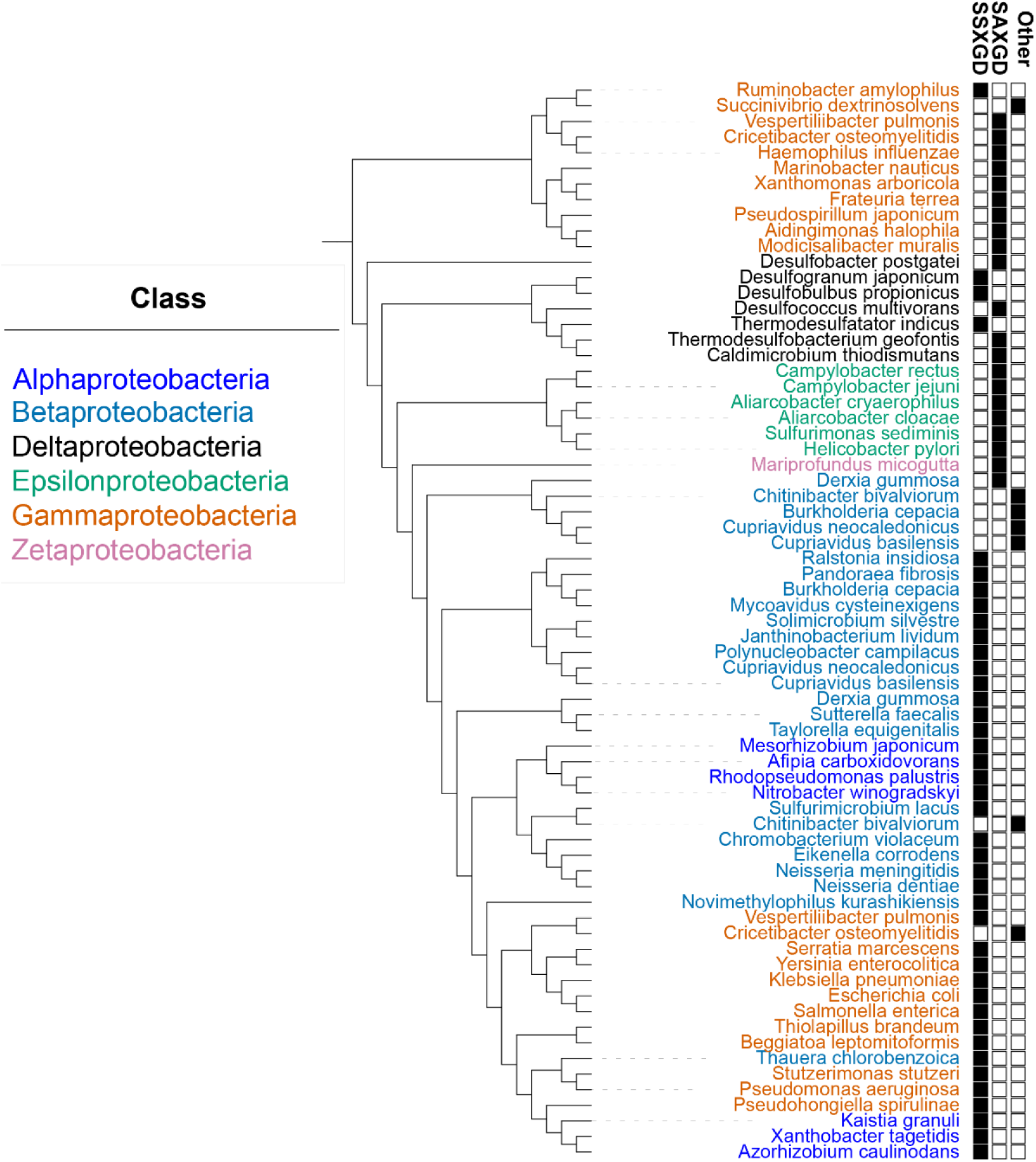
Phylogenetic tree of HepI sequences in proteobacteria.

We hypothesize that the two major alleles of the substrate-binding motif present in γ proteobacteria may contribute to the structural variability in lipid A and the heterogeneity of lipopolysaccharide biosynthesis within the clade. To test this, we compared the wild-type HepI structure from *E. coli* (structure available in PDB) to an *in silico* mutant structure with the *H. influenzae* binding motif.

### 3.2 Overall Trajectory and Protein Dynamics Differences between HepI variants

The HepI *E. coli* protein structure (wt-HepI) and the *H. influenzae*-like structural model (mt-HepI) were subjected to 100 ps NVT and 100 ps NPT equilibration prior to a 1 μs molecular dynamics (MD) simulation. The two simulations exhibited large global structural changes, that were largely the result of structural variations in the linker region for wt-HepI. Unsurprisingly, the mt-HepI simulation took longer to establish an equilibrium structure, due to the need to rearrange to adapt to the introduction of amino acid substitutions, but both proteins showed minimal variation for the 700 ns – 1 us portion of the simulation, and therefore the remaining analyses focused on observations over this time range.

The backbone root mean square deviation (RMSD) of the last 300 ns of the trajectory for wt-HepI and mt-HepI was calculated with the unchanged C-terminal used for generating the structural superposition. Structural snapshots of the protein trajectories for the wt- and mt-HepI variants were generated at 700, 800, 900 and 1000 nanoseconds to examine the structural variations observed over these time periods. RMSD for the pairwise comparisons of structures at the various time points for either the wt-HepI or mt-HepI were less than 3.0 Å (Figure S3 & Table S2), with wt-HepI having structural RMSDs of 2.931, 1.604 and 1.695 and mt-HepI having RMSD values of 2.438, 2.112 and 1.775 for comparison of the pairwise snapshots of 700 ns to 800 ns, 800 ns to 900 ns and 900 ns to 1000 ns, respectively. However, the comparisons between wt-HepI and the mt-HepI at any given timepoint were noticeably more diverged from each other, with major differences in the angle of the opening between the N- and C-terminal Rossman-like domains and the positioning of loops (Figure 4 & Table S3) resulting in an average deviation of ∼4.2 for each paired timepoint. The Cα root mean square fluctuations (CαRMSF) for wt-HepI reveal residues with the greatest fluctuations in 60s, 120s, 230s and 290s, consistent with prior simulations [21, 22]. The CαRMSF for mt-HepI exhibited fluctuations in all of these regions, but to a lesser extent than for the wt-HepI simulation, except in the linker region and in various parts of the C-terminal domain (Figure S4).

**Figure 4:**
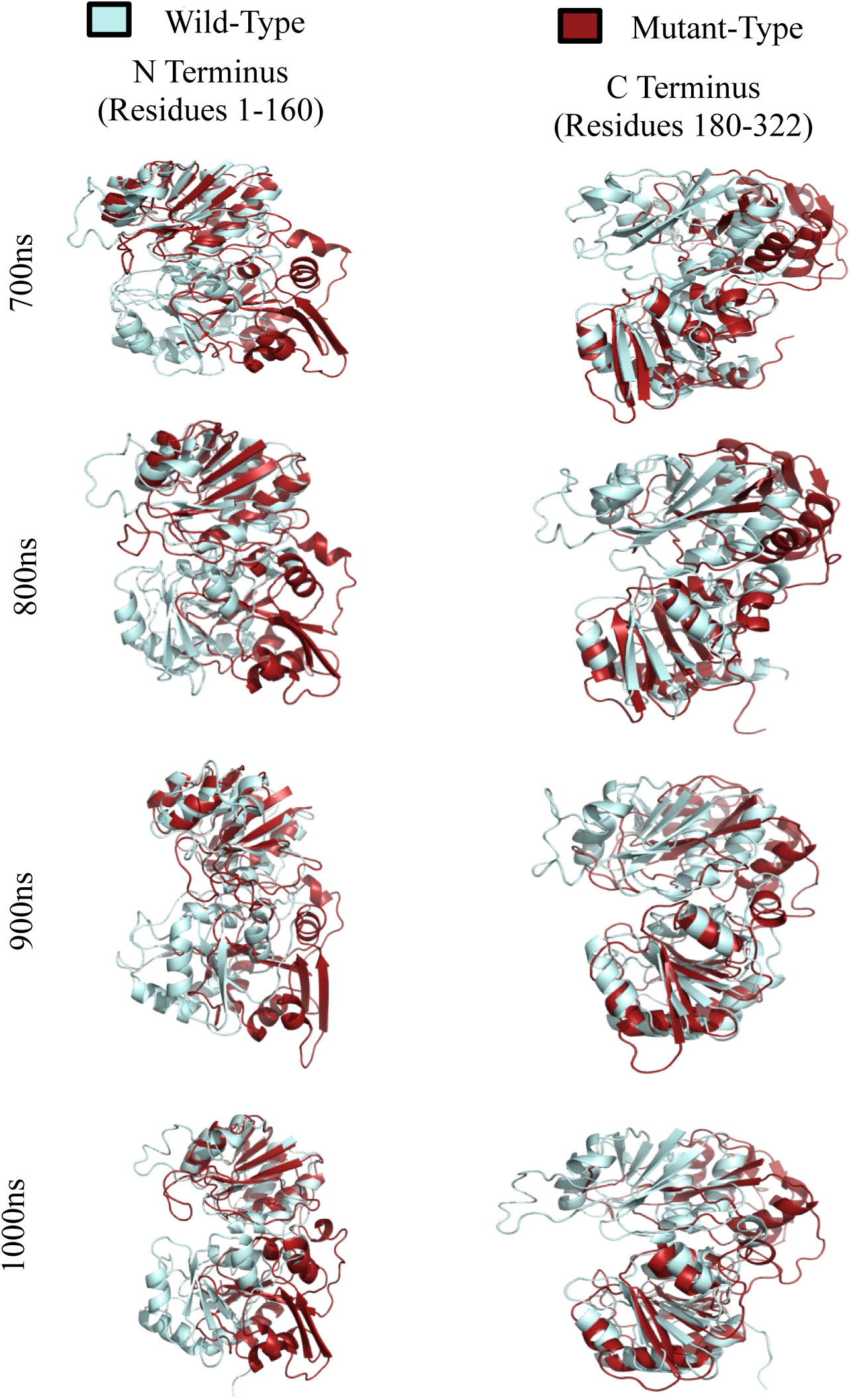
Aligned Superpositions of wt-HepI (light blue) and mt-HepI (maroon) over intervals of 100 nanoseconds, matched by N (left) and C (right) Terminus domains. Generated in PyMOL software.

### 3.3 Reduction in Binding Pocket Area and Volume due to mutation of HepI

The CASTp analysis of the binding pocket area and volume over the converged trajectory intervals provided crucial insights into how mutations could alter the structural and functional dynamics of HepI. Our analysis focused on the binding pockets of the N-terminal Lipid A binding domain. Prior computational and experimental studies have revealed the amino acid residues in the 10-13, 60-63, and 120 regions as critical for the enzyme’s function [21, 24–26]. For each structure, we recorded the area and volume of the binding pocket (Table 1) encompassed by these regions. This data collection allowed us to capture the dynamic variations in pocket dimensions across the sampled time points. By averaging the area and volume measurements from the sampled structures, we obtained a comprehensive view of the binding pocket dimensions for both the wildtype and mutant HepI proteins. Our analysis revealed a significant reduction in the binding pocket area and volume in the mutant HepI compared to the wildtype. This decrease was likely a result of mutation-induced structural rearrangements, leading to a more compact and stable protein conformation.

**Table 1:** Binding pocket area and volume over the converged trajectory.

| **System** | **Area (Å<sup>2</sup>)** | **Volume (Å<sup>3</sup>)** |
| --- | --- | --- |
| wt-700ns.pdb | 1041.034 | 904.512 |
| wt-800ns.pdb | 285.262 | 361.926 |
| wt-900ns.pdb | 330.034 | 466.292 |
| wt-1000ns.pdb | 1118.078 | 1789.968 |
| mt-700ns.pdb | 158.028 | 167.907 |
| mt-800ns.pdb | 327.739 | 207.690 |
| mt-900ns.pdb | 308.910 | 168.637 |
| mt-1000ns.pdb | 683.153 | 891.172 |

### 3.4 Minimum Distance from Center of Mass between Active Sites Decreased

The minimum distance from the center of mass between active sites within regions 10-13, 60-63, and 120 decreased in the mt-HepI compared to the wt-HepI. This observation suggested the mutation likely enhanced interactions between residues within and between these regions (Figure 5a-c). These interactions bring active site residues closer together, reducing the overall distance between their respective centers of mass. The closer proximity of active site residues reflects a more compact and stabilized protein structure. The mutation appeared to enhance the interactions between residues within and between the critical active site regions. By bringing these residues closer together, the protein adapted to stabilize its structure despite the potential destabilizing effects of the mutation.

**Figure 5:**
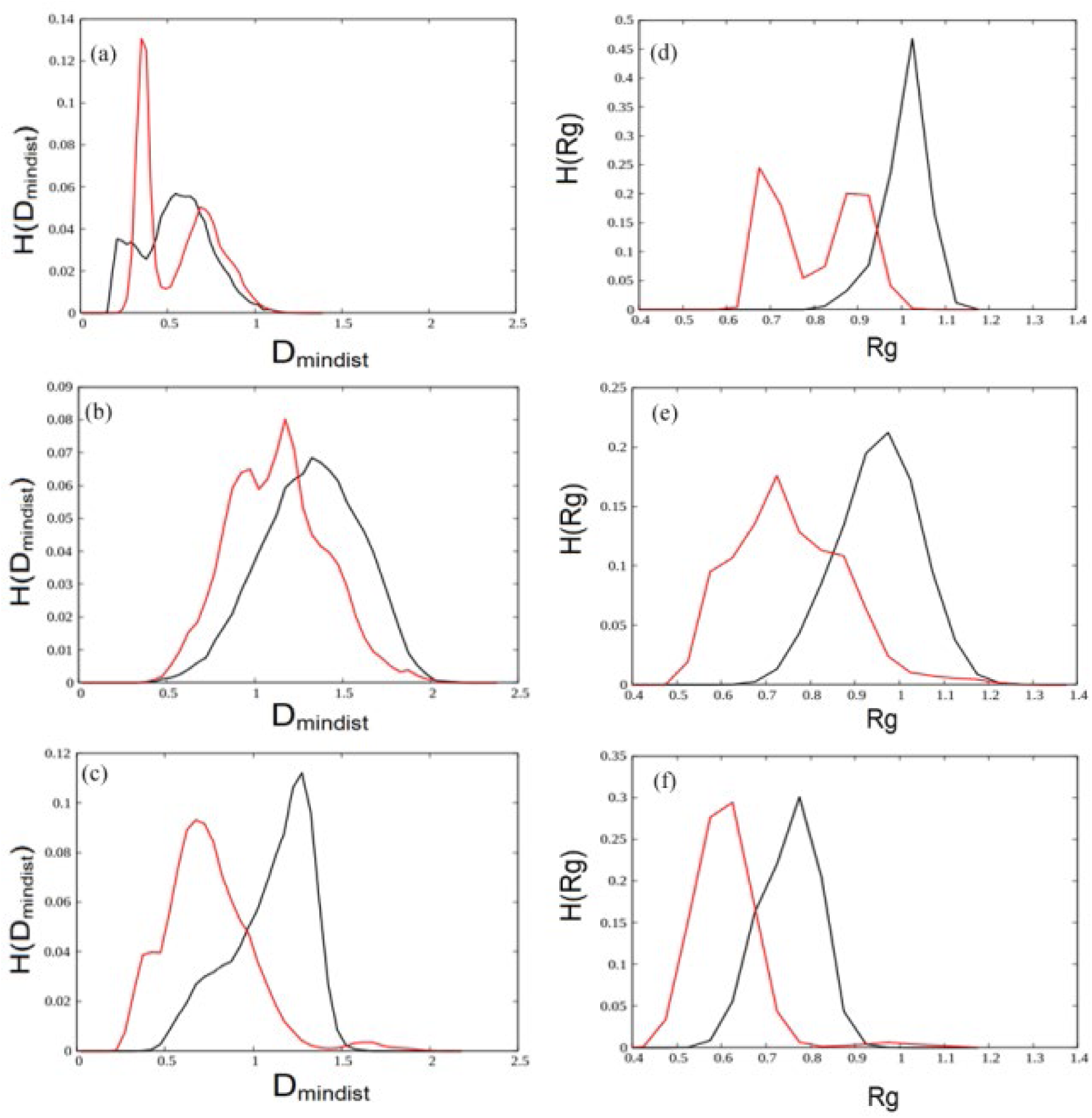
Histogram distribution of the minimum distance from the center of mass between active sites residues (a) S10A and R60V, (b) R60V and R120 (c) S10A and R120 in wt-HepI (black line) and mt-HepI (red line). Histogram distribution of the radius of gyration (Rg) between wt-HepI (black line) and mt-HepI (red line) in (d) S10A and R60V (e) R60V and R120 (f) S10A and R120

### 3.5 Comparative Analysis of Radius of Gyration (Rg) between Variants of HepI

The radius of gyration (Rg) could be defined by the overall dimension of the protein in a dynamic system and was derived from computing the mass-weighted root mean square deviation of all atoms from the center of mass. In conformational analysis, Rg represented the moment of inertia of a group of atoms relative to their center of mass. Due to the multidomain GT-B structure of HepI, a comparative analysis of the radius of gyration between wt-HepI and mt-HepI, focusing on the N-terminal regions encompassed by amino acid residues 10-13, 60-63, and 120 (Figure 5d-f).

In mt-HepI, Rg was reduced compared to wt-HepI. This reduction indicated that the protein adopts a more compact conformation, which likely reduced the volume of the binding pocket. Such compaction might stabilize the structure in response to the mutation, leading to the mutated HepI being more tightly packed. This tighter packing likely serves as a stabilizing response to the mutation, which might otherwise destabilize the protein. By reducing the Rg, the protein increased its local stability at the expense of flexibility (as described above). Additionally, the reduced Rg might reflect a functional adaptation to the mutation, resulting in a smaller, more rigid active site. This could enhance substrate selectivity or alter catalytic efficiency, compensating for the structural changes induced by the mutation.

### 3.6 Analysis of Solvent Accessible Surface Area (SASA) in Wildtype and Mutant Variants of HepI

Measurement of SASA provided insight into the compactness of the hydrophobic core, a key factor in biomolecular stability. The alteration of solvent accessible surface area (SASA) between the wt-HepI and mt-HepI over simulation time was depicted in Figure 6(a-c). The SASA of the mt-HepI protein was significantly decreased compared to the wt-HepI, particularly in the regions between residues 10-13 to 60-63 and 60-63 to 120, with a smaller decrease observed in the 10-13 to 120 region. The mutation appears to have caused more surface area of the protein to be buried, reducing its exposure to the solvent. This burying effect was attributed to the tighter packing of residues, which reduced the accessibility of certain regions within the protein structure. Conformational changes induced by the mutation alter the protein’s surface topology, leading to a decrease in the accessible surface area. This alteration reflected a structural adaptation necessary for maintaining enzymatic activity in the mutated protein variant.

**Figure 6:**
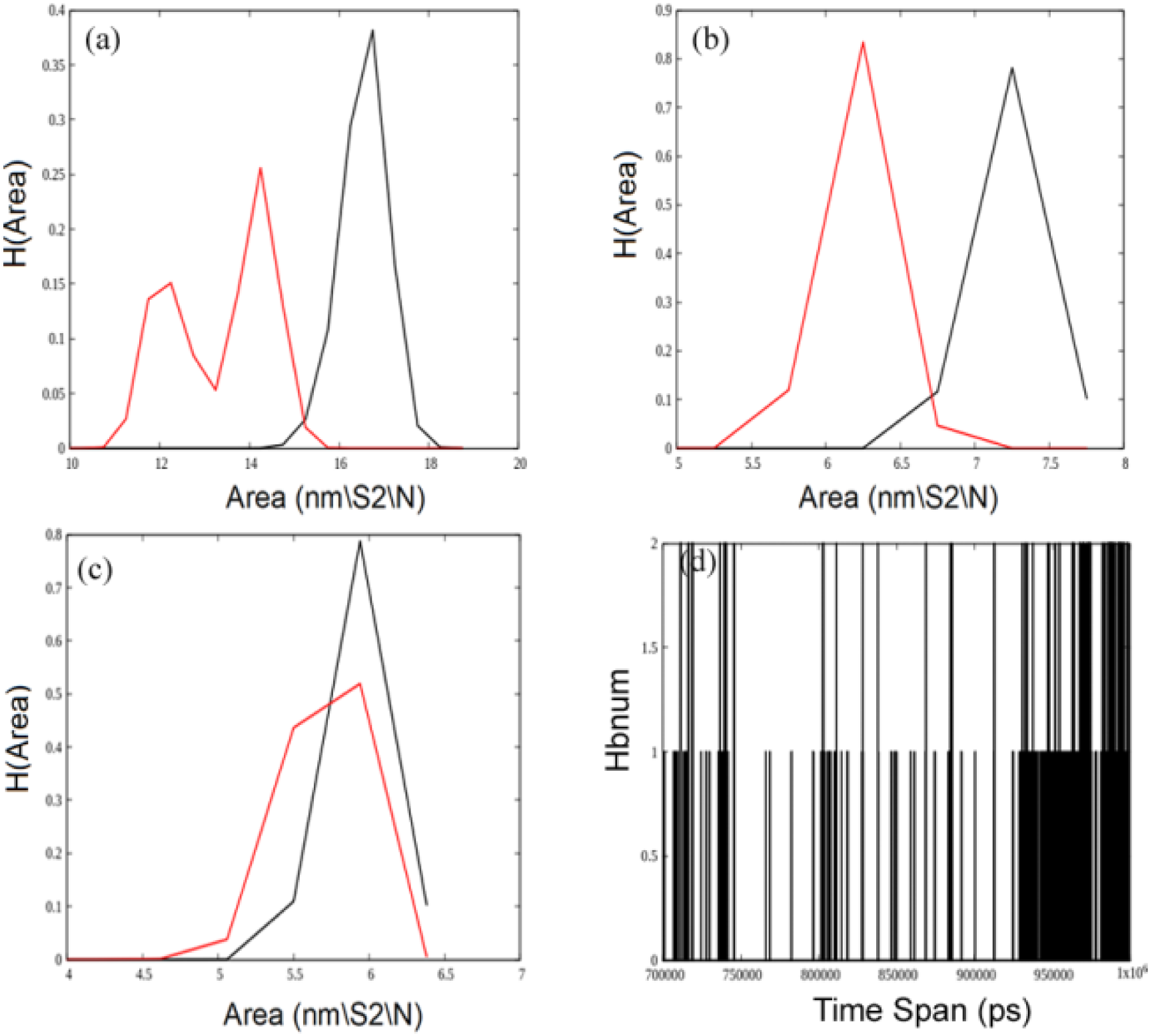
Histogram distribution of solvent accessible surface area (SASA) between wt-HepI (black line) and mt-HepI (red line) in (a) S10A and R60V, (b) R60V and R120 (c) S10A and R120. (d) Histogram plot of the number of hydrogen bond between S10 and R60 in wt-HepI (black line).

### 3.7 Impact of S10-R60 Hydrogen Bond Disruption on HepI Structure and Function

The mutation in HepI disrupted the hydrogen bond between S10 and R60 regions, leading to significant conformational shifts (Figure 6d). The regions previously stabilized by this bond might reposition themselves, altering the overall structure. Such changes could disrupt the proper folding of HepI, affecting its shape and how it interacts with substrates and other molecules. Given the proximity of S10 and R60 to the active site, the loss of their interaction can directly impact the active site’s structure. This disruption can alter the binding affinity and orientation of substrates, potentially reducing the catalytic efficiency of HepI. The hydrogen bonding network for the wild-type (wt-HepI) and mutant (mt-HepI) structures was analyzed at various time points from the converged trajectory (Figure S5). Each position in the HepI protein is represented by a point on the circle’s perimeter, with lines connecting two points indicating the presence of a hydrogen bond. The color of the line distinguishes between bonds present in wt-HepI (black), mt-HepI (red), or both (yellow). This analysis reveals that the wt-HepI structure exhibits a higher number of hydrogen bond interactions compared to the mt-HepI. Understanding the impacts of this disruption provided valuable insights into how mutations influence protein function, highlighting the delicate balance between protein structure and function.

### 3.8 Structural Insights from Secondary Structure Analysis of wildtype and mutant HepI

The structural consequences of a mutation in HepI revealed significant insights through a detailed secondary structure analysis using DSSP (Figure S6). This analysis indicated an increase in β sheets, β bridges, and bends in the mt-HepI compared to the wt-HepI. The wt-HepI showed a predominance of helical portions, including α helices and 3_10 helices, which was less prominent in the mutated form (Figure 7a-c). The increase in β sheets and β bridges in the mt-HepI might be a compensatory mechanism to stabilize the protein structure after the mutation disrupted other stabilizing interactions, such as the S10-R60 hydrogen bond (Figure 7d-f). Β sheets and bridges were known for their ability to provide rigidity and structural integrity due to extensive hydrogen bonding networks between backbone atoms. Β sheets formed through hydrogen bonds between the backbone amides and carbonyls of different strands. The mutation might induce a rearrangement that favors the formation of these additional hydrogen bonds, thereby increasing the presence of β sheets and bridges. The mutation could cause a reorganization of secondary structure elements, where regions that were previously in other conformations (e.g., α helices) may shift to form β sheets and bridges. This shift could be a response to the need for maintaining a stable core structure despite the destabilizing mutation. An increase in bends might indicate an adaptive increase in flexibility in certain regions of the protein. This could help HepI maintain its functional dynamics, allowing it to adjust its conformation to accommodate substrates despite the mutation-induced structural changes.

**Figure 7:**
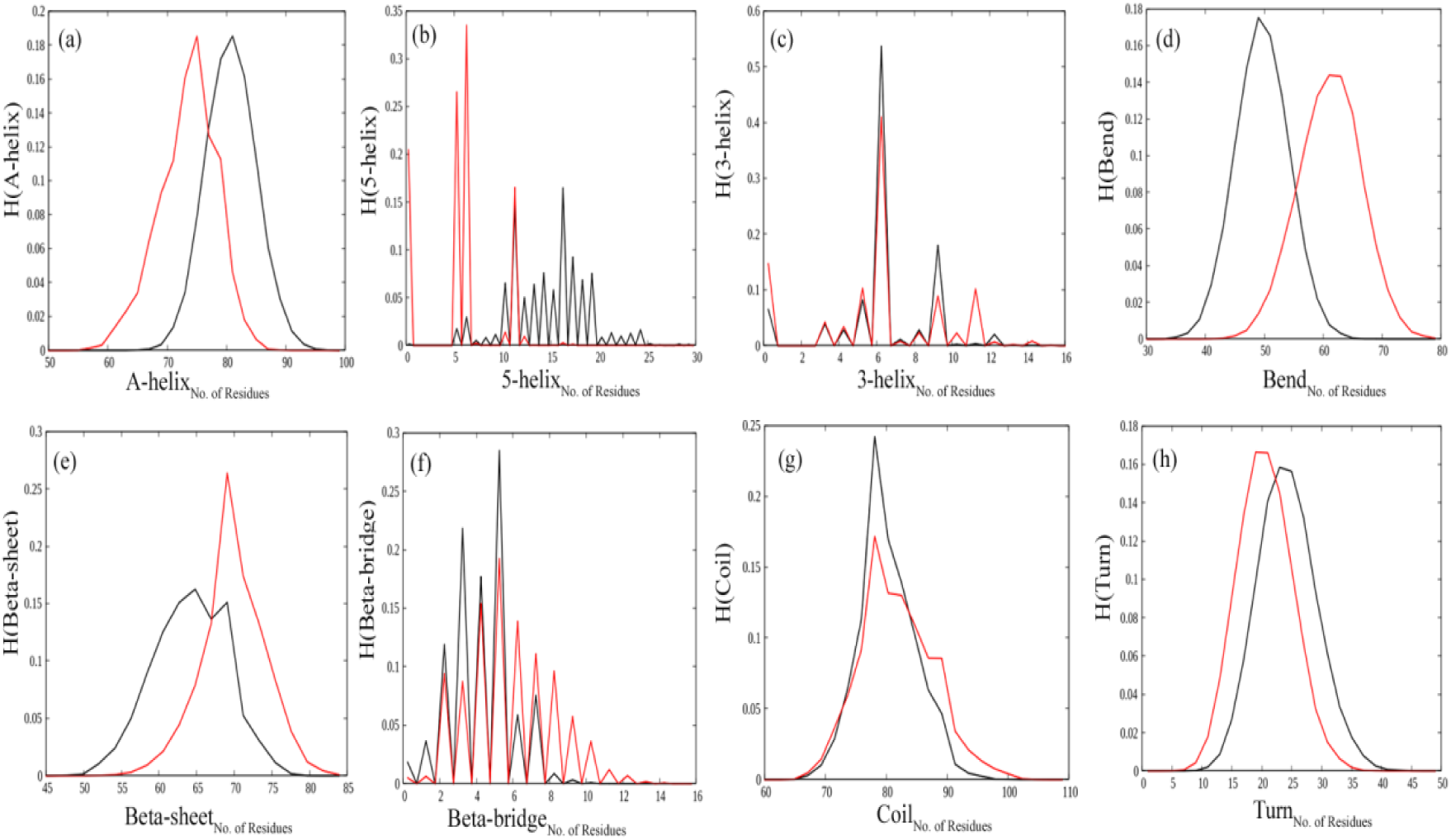
Histogram distribution of the comparative secondary structures between wt-HepI (black line) and mt-HepI (red line) (a) A-helix (b) 5-helix (c) 3-helix (d) Bend (e) beta sheet (f) beta bridge (g) coil (h) Turn

### 3.9 Principal Component Analysis (PCA) Reveals Dynamic Differences Between Wildtype and Mutant HepI

Essential dynamics, specifically Principal Component Analysis (PCA), was utilized to gain a comprehensive perspective on the dynamic properties observed in MD simulations. The projection of the first two eigenvectors (ev1 vs ev2) was conducted for both wt-HepI and mt-HepI proteins (Figure S7). It was observed that the first few principal components (PCs) captured the majority of the protein’s movement, making them reliable for representing its motion, particularly PC1. Figure 8a-b depicted a PCA scatter plot for wt-HepI and mt-HepI, highlighting significant differences between the two systems. This distinction was evident from the characteristic structures plotted along the directions defined by the first two principal components. The scatter plot clearly illustrated that the eigenvectors computed from the MD trajectories for each system varied noticeably, indicating substantial differences in protein motion between wt-HepI and mt-HepI.

**Figure 8:**
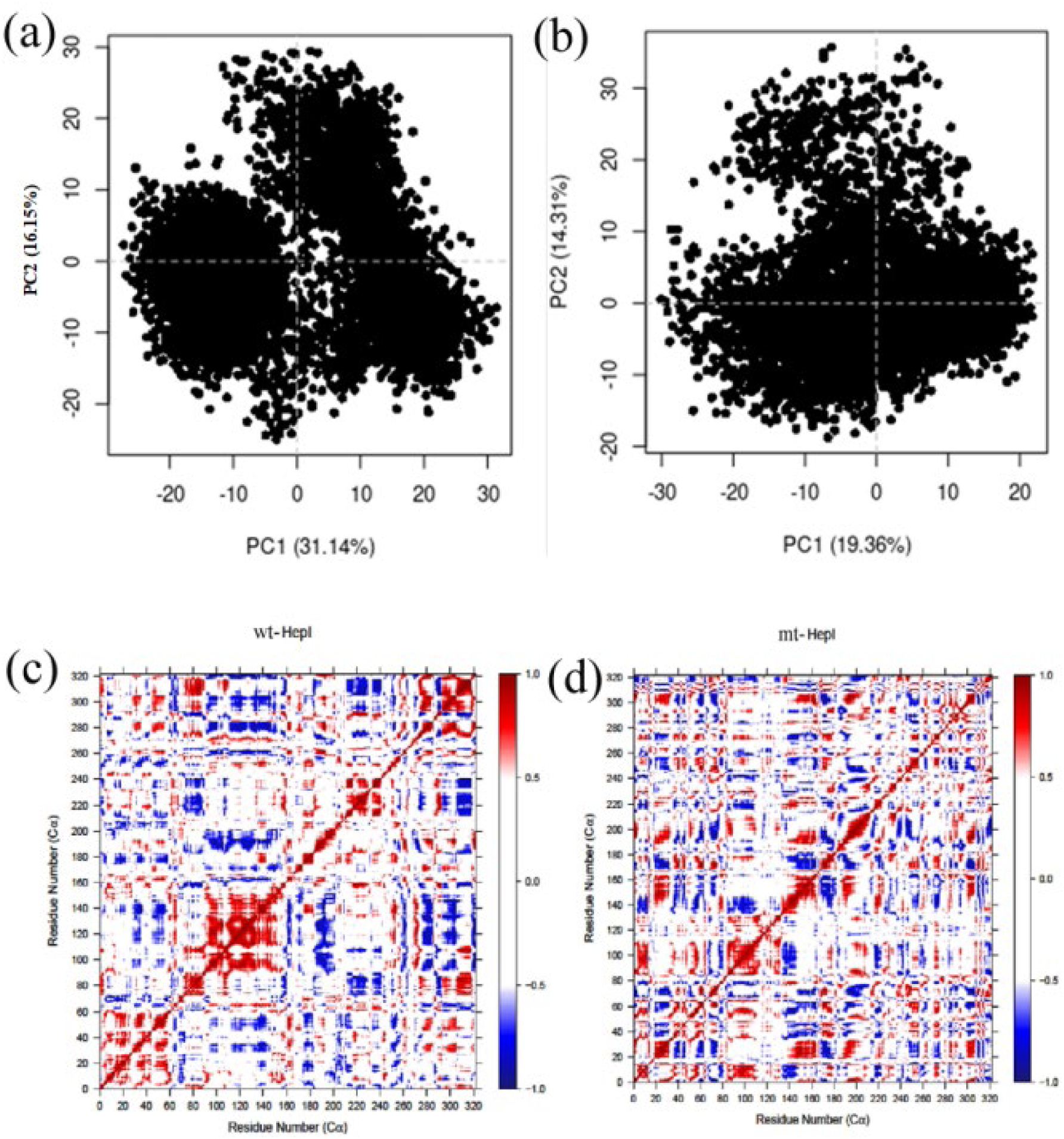
PCA scatter plot on Cα atoms along first two principal components, PC1 and PC2 respectively showing difference in motion in (a) wt-HepI and (b) mt-HepI. Cross-correlation maps reflecting relative motions between residues of (c) wt-HepI and (d) mt-HepI.

In wt-HepI, the PCA analysis revealed a bifurcation of the PC1 vs. PC2 curve into two distinct clusters, indicating the presence of multiple dominant conformational states. This observation suggests that the wildtype protein exhibits a higher degree of conformational flexibility and complexity. Conversely, mt-HepI showed a single scatter plot, indicating a more uniform conformational distribution and less flexibility. This suggests that the mutation in mt-HepI results in a protein that adopts a more restricted set of conformations, reflecting reduced dynamism. Furthermore, the analysis demonstrated that wt-HepI has a higher proportion of variance associated with the first few eigenvalues compared to mt-HepI. This higher variance indicates that the primary modes of motion in wt-HepI capture a larger portion of the total movement within the protein, highlighting its greater conformational diversity. In contrast, mt-HepI exhibited a lower proportion of variance in the top eigenvalues, suggesting that the mutation has constrained the protein’s conformational space. This resulted in a more rigid and less dynamic structure compared to the wildtype protein.

### 3.10 Dynamic Cross-Correlation Matrix (DCCM) Analysis Reveals Mutation-Induced Residue Interactions in HepI Variants

The dynamic interplay within HepI enzymes, especially wt-HepI and mt-HepI, representing *E. coli* and *H influenza*, was not just a tale of molecular movements but also one of structural adaptation and changes in the extent of correlated atomic fluctuation networks. The observed changes in DCCM analysis revealed significant shifts in residue interactions and correlation patterns due to mutation, with mt-HepI introducing profound shift in this dynamic landscape. (Figure 8c-d). Whereas the wt-HepI has extensive collections of nearby residues that demonstrate correlated atomic fluctuations (with positively correlated motions being depicted in red, and negatively correlated mothions being depicted in blue), the wt-HepI shows much smaller collections of residues that have correlated motions. Specifically, in the wt-HepI residues 0-60, 70-160, 180-260 and 290-330 show significant amounts of positively correlated atomic fluctuations, whereas the correlated motions observed in mt-HepI are each isolated to a given secondary structural feature. Examination of the difference in the motions of these two DCC matrixes, shows that very few regions maintain their pattern of correlated motions (depicted by red or blue coloring), while numerous regions now have correlation patterns that are the opposite in the two simulations (magenta) (Figure S8). As has been previously observed the wt-HepI, residues in the 10s and 60s move in concert, as do the 60s and 120s regions allowing for these regions to move in concert to bind the substrate. In the mt-HepI, while these regions are still exhibiting coordinated motions, the overall number of residues in each region and magnitude of the correlation are reduced. The decrease in size of regions exhibiting correlated motions in mt-HepI suggests that the mutation has perhaps disrupted longer range interactions as a direct consequence of structural rearrangements within the protein, as the mutation alters the protein’s internal landscape.

## 4. Discussion

The structural heterogeneity observed in lipopolysaccharide (LPS) biosynthesis among proteobacteria reflects the sensitivity to heptosyltransferase sequence motifs to environmental pressures, influencing lipid A structural variations such as hexosamine type, phosphorylation degree, and acyl chain characteristics. γ proteobacteria occupied an evolutionary niche bridging other proteobacterial groups, highlighting a convergence of sequence motifs and evolutionary traits. This positioning might stem from shared ecological roles or adaptive pressures across proteobacterial classes.

The phylogenetic divergence within γ proteobacteria, particularly in the orders Enterobacteriales and Pseudomonales, underscores their complex evolutionary history. These groups display a blend of sequence motifs similar to those found in α, β, ε, δ, and ζ proteobacteria, indicating adaptive strategies that balance structural integrity with functional versatility. Phylogenetic analysis reveals that γ proteobacteria exhibit greater flexibility in their genetic sequences and functional capabilities compared to other bacterial classes. This adaptability allows them to undergo a wide range of genetic variations while maintaining functionality. Such genetic flexibility likely contributes to their ability to thrive in diverse environments and adapt to various ecological niches, highlighting their evolutionary resilience and versatility.

For α, β, and part of the γ proteobacteria, including *E. coli*, α and β proteobacteria exhibited the distinctive SSXGD conserved substrate binding motif at amino acid positions 9-13, along with the RRWRK conserved region at positions 60-64 and utilize Kdo2-lipid A as the N-terminal binding sugar acceptor substrate. ε, δ, ζ and the remaining γ proteobacteria, including *H. influenzae*, typically showed a S10A mutation, resulting in the SAXGD motif, followed by the VVYDK/IVYDK conserved region at positions 60-64. It is known that the Kdo transferase for *H. influenzae* HepI utilizes exclusively Kdo-phosphate-lipid A as its N-terminal sugar acceptor substrate and not Kdo2-lipid A.

The structural heterogeneity in lipopolysaccharide (LPS) biosynthesis among proteobacteria is significantly influenced by the presence and function of their heptosyltransferases (and the Kdo-transferase which attaches the preceding sugars onto LipidA), contributing to the diverse LPS structures observed across different bacterial species. In *wild-type* bacteria regardless of whether the LPS is harboring a Kdo2 or Kdo-phosphate functionalized Lipid A, a heptose is attached at position 5 of the Kdo. Two types of heptosyltransferase have evolved in response to these different lipid A acceptors, such that Kdo2-lipid A and Kdo-phosphate-lipid A, require distinct heptosyltransferases to accommodate the the varying sugar acceptor sizes. This research is the first study to identify the specific sequence variations that result in this difference.

Researchers have previously dubbed the Kdo-phosphate utilizing Heptosyltransferase from *H. influenzae* as “OpsX-like” [27], in reference to the gene having similarity to the opsX gene encoding a heptosyltransferase in *Xanthomonas campestris*. Investigation of the LPS structure of different strains of *X. campestris* revealed that while the B100 strain has heptoses appended onto the Kdo-phosphate-lipid A [28], many *Xanthomonas* strains do not append a heptose sugar onto Kdo in its LPS [29, 30]. Analysis of recombinant *E. coli* strains *in vivo* [31] and *in vitro* kinetic analyses of the *E. coli* HepI [32] have previously shown that the enzyme can accept the smaller Kdo-phosphate, but with lower enzymatic turnover. However, *H. influenzae* “OpsX-like” heptosyltransferase cannot transfer heptose onto a Kdo disaccharide acceptor, suggesting that the larger Kdo side chain does not fit into the binding pocket of “OpsX-like” heptosyltransferases [27]. This supports the hypothesis that structural adaptations in heptosyltransferases allow for specific interactions with their substrates. Of interest is that the opsX gene has long been described as a virulence factor [33], and that it contribute to bacterial resistance to antimicrobial peptides [33]. Preliminary analyses has shown that some organisms contain both “OpsX-like” and other HepI encoding genes, and further exploration of the differences between these different heptosyltransferases are ongoing.

The observation that heptosyltransferase I sequence motifs allow for the prediction of LPS structural features, reflecting the sequence-structural adaptations of HepI enzymes across proteobacteria could be useful in future analyses of LPS variability. These adaptations highlight their evolutionary resilience and functional versatility, contributing to the diverse LPS structures observed among different bacterial species. The structural heterogeneity in LPS biosynthesis among proteobacteria reflects the sensitivity of HepI sequence motifs to environmental pressures, influencing lipid A structural variations such as hexosamine type, phosphorylation degree, and acyl chain characteristics [34, 35]. γ-proteobacteria, in particular, occupy an evolutionary niche bridging other proteobacterial groups, highlighting a convergence of sequence motifs and evolutionary traits. This positioning might stem from shared ecological roles or adaptive pressures across proteobacterial classes.

The phylogenetic divergence within γ-proteobacteria, particularly in the orders Enterobacteriales and Pseudomonales, underscores their complex evolutionary history. These groups display a blend of sequence motifs similar to those found in α, β, ε, δ, and ζ proteobacteria, indicating adaptive strategies that balance structural integrity with functional versatility. This reveals that γ-proteobacteria exhibit greater flexibility in their genetic sequences and functional capabilities compared to other bacterial classes. This adaptability allows them to undergo a wide range of genetic variations while maintaining functionality, contributing to their ability to thrive in diverse environments and adapt to various ecological niches [36, 37].

Since HepI variants from these two organisms perform chemistry on tetrasaccharide or trisaccharide glycolipids, respectively, we hypothesized that the active site geometry, volume, and dynamics would be noticeably altered in response to the substrate differences. These studies demonstrated that these sequence changes result in significant shifts in the binding pocket’s volume, influencing how these enzymes accommodate their substrates. These mutations also induced structural rearrangements in HepI, evidenced by changes in binding pocket volume, radius of gyration, and residue interactions, reflecting adaptive responses to maintain enzymatic activity. Anti-correlated movements between residues and shifts in secondary structure elements indicated compensatory mechanisms to stabilize the protein amidst mutation-induced perturbations, ensuring the enzyme retained functionality by enhancing substrate selectivity and catalytic efficiency despite altered structural dynamics.

Furthermore, reductions in solvent-accessible surface area (SASA) and increases in β structures (sheets, bridges, bends) in mt-HepI highlighted additional stabilization mechanisms. These changes facilitated tighter packing and enhanced hydrogen bonding, maintaining structural integrity and enzymatic function under mutation-induced stress. The adaptation of HepI to genetic variations underscored its capability to sustain enzymatic activity and stability, crucial for its biological roles in diverse environmental contexts. Overall, these findings illustrate how the sequence-structural adaptations of HepI enzymes across proteobacteria reveal their evolutionary resilience and functional versatility. The diverse structural features and functional capabilities of these enzymes allow proteobacteria to thrive in various environments and adapt to ecological pressures, underscoring the evolutionary success of these bacterial groups.

## 5. Conclusions

(The conclusion section should present the main conclusions of your study. You may have a stand-alone conclusions section or include your conclusions in a subsection of your discussion or results and discussion section.)

In response to different environmental context and selective pressures, Gram-negative bacteria have demonstrated the ability to alter the number of Kdo residues incorporated into their LPS structure, and sequence analysis of HepI which adds a heptose residue onto Kdo can help reveal whether an organism incorporates one or two Kdos into its structure. These adaptations correspond to intricate adjustments in active site residues, balancing structural stability with functional integrity in response to genetic mutations, ultimately impacting the structural and functional integrity in their LPS biosynthesis pathways.

In summary, understanding the structural adaptations of HepI enzymes across proteobacteria sheds light on their evolutionary resilience and functional versatility. These enzymes exemplify nature’s ability to fine-tune molecular machinery in response to environmental pressures, ensuring bacterial survival and adaptation in diverse ecological niches. This study explored how sequence diversity and structural adaptations in HepI enzymes underpinned the dynamic interplay between protein structure and function, essential for deciphering their roles in bacterial physiology and pathogenesis. Understanding the functional implications of HepI adaptations in bacterial pathogenesis is crucial. Studies focusing on the correlation between specific HepI motifs and bacterial virulence could inform vaccine development and therapeutic interventions targeting LPS-mediated immune responses. These subtle sequence differences in HepI lead to significant alterations in its structural dynamics and functional properties, potentially affecting multiple biological processes in these organisms.

## 6. Acknowledgements

The authors gratefully acknowledge the use of the CRAY computing facility at the Technical Research Centre, S. N. Bose National Centre for Basic Sciences, for providing computational resources crucial to this research.

## 7. Author contributions

A.M.G, P.A. and E.A.T were all involved in the Conceptualization, Data curation, Formal analysis, Methodology, Validation, Visualization and Writing – review and editing; A.M.G. was responsible for the Investigation and Writing – original draft; B.A.H. assisted with Methodology and Visualization; J.B. assisted with the Visualization; P.A. and E.A.T were involved in the Project administration and Supervision

## 8. Funding sources

This research did not receive any specific grant from funding agencies in the public, commercial, or not-for-profit sectors.

## Notes

### Competing Interest Statement

The authors have declared no competing interest.

