## Supplemental Information for "Structural Adaptations of HepI Enzymes in Proteobacteria: Insights into Evolutionary Resilience and Functional Dynamics"

Table S1: List of Organisms used in the study.

| **Species** | **Gene Id** |
|---|---|
| Haemophilus influenzae | NZ_LN831035.1_784 |
| Cricetibacter osteomyelitidis | NZ_SLYB01000004.1_69 |
| Cricetibacter osteomyelitidis | NZ_SLYB01000004.1_2 |
| Vespertiliibacter pulmonis | NZ_CP016615.1_1409 |
| Vespertiliibacter pulmonis | NZ_CP016615.1_655 |
| Ruminobacter amylophilus | NZ_FOXF01000026.1_10 |
| Succinivibrio dextrinosolvens | NZ_FUXX01000002.1_7 |
| Frateuria_A terrea | NZ_FOXL01000005.1_51 |
| Xanthomonas arboricola | NZ_JZEF01000003.1_242 |
| Marinobacter nauticus | NZ_RBJB01000006.1_107 |
| Pseudospirillum japonicum | NZ_FNYH01000015.1_33 |
| Modicisalibacter muralis | NZ_FNGI01000004.1_238 |
| Aidingimonas halophila | NZ_FNNI01000003.1_271 |
| Novimethylophilus kurashikiensis | NZ_BDOQ01000003.1_221 |
| Chromobacterium violaceum | NC_005085.1_216 |
| Eikenella corrodens | NZ_LT906482.1_933 |
| Neisseria meningitidis | NZ_LR134525.1_1817 |
| Neisseria dentiae | NZ_CP059570.1_2209 |
| Beggiatoa leptomitoformis | NZ_CP012373.2_2719 |
| Salmonella enterica | NC_003197.2_3666 |
| Escherichia coli | NZ_CP033092.2_84 |
| Klebsiella pneumoniae | NZ_KN046818.1_4465 |
| Yersinia enterocolitica | NZ_LR590469.1_3154 |
| Serratia marcescens | NZ_CP071238.1_4598 |
| Thauera chlorobenzoica | NZ_CP018839.1_1208 |
| Pseudomonas aeruginosa | NZ_LN831024.1_5211 |
| Stutzerimonas stutzeri | NC_015740.1_3782 |
| Pseudohongiella spirulinae | NZ_CP013189.1_305 |
| Thiolapillus brandeum | NZ_AP012273.1_2883 |
| Chitinibacter bivalviorum | NZ_CP058627.1_2340 |
| Chitinibacter bivalviorum | NZ_CP058627.1_3135 |
| Sulfurimicrobium lacus | NZ_AP022853.1_104 |
| Derxia gummosa | NZ_AXWS01000007.1_159 |
| Derxia gummosa | NZ_AXWS01000007.1_843 |
| Sutterella faecalis | NZ_CP040882.1_1405 |
| Taylorella equigenitalis | NC_018108.1_52 |
| Janthinobacterium lividum | NZ_UGJH01000001.1_4964 |
| Solimicrobium silvestre | NZ_PUGF01000001.1_118 |
| Cupriavidus basilensis | NZ_CP062803.1_2551 |
| Cupriavidus basilensis | NZ_CP062803.1_3129 |

| | |
|---|---|
| Polynucleobacter campilacus | NZ_NGUP01000003.1_882 |
| Ralstonia insidiosa | NZ_VZPV01000001.1_1237 |
| Pandoraea fibrosis | NZ_CP047385.1_1414 |
| Mycoavidus cysteinexigens | NZ_AP018150.1_1824 |
| Burkholderia cepacia | NZ_CP012981.1_1650 |
| Burkholderia cepacia | NZ_CP012981.1_1730 |
| Cupriavidus neocaledonicus | NZ_AQUR01000101.1_82 |
| Cupriavidus neocaledonicus | NZ_AQUR01000105.1_270 |
| Mariprofundus micogutta | NZ_BDFD01000009.1_31 |
| Helicobacter pylori | NZ_LS483488.1_1460 |
| Campylobacter_A rectus | NZ_CP012543.1_1620 |
| Sulfurimonas sediminis | NZ_CP041235.1_1963 |
| Campylobacter_D jejuni | NZ_LN831025.1_1213 |
| Aliarcobacter cloacae | NZ_CP053833.1_2073 |
| Aliarcobacter cryaerophilus | NZ_CP032823.1_512 |
| Caldimicrobium thiodismutans | NZ_AP014945.1_151 |
| Desulfococcus multivorans | NZ_CP015381.1_2967 |
| Afipia carboxidovorans | NC_015684.1_2676 |
| Azorhizobium caulinodans | NC_009937.1_2316 |
| Desulfobacter postgatei | NZ_CM001488.1_1672 |
| Desulfobulbus propionicus | NC_014972.1_2805 |
| Desulfogranum japonicum | NZ_AUCV01000001.1_312 |
| Kaistia granuli | NZ_AQYH01000003.1_260 |
| Mesorhizobium japonicum | NC_002678.2_2018 |
| Nitrobacter winogradskyi | NC_007406.1_1086 |
| Rhodopseudomonas palustris_J | NC_007778.1_1604 |
| Thermodesulfatator indicus | NC_015681.1_1320 |
| Thermodesulfobacterium geofontis | NC_015682.1_1164 |
| Xanthobacter tagetidis | NZ_RCTF01000022.1_17 |

Table S2: Pairwise comparisons of evolution of structures at various time points from the converged trajectory for wt-HepI and mt-HepI

| wt-HepI | 700 | 800 | 900 | 1000 |
|---|---|---|---|---|
| 700 | x | 2.931 | | |
| 800 | 2.931 | x | 1.604 | |
| 900 | | 1.604 | x | 1.695 |
| 1000 | | | 1.695 | x |
| mt-HepI | 700 | 800 | 900 | 1000 |
| 700 | x | 2.438 | | |
| 800 | 2.438 | x | 2.112 | |
| 900 | | 2.112 | x | 1.775 |
| 1000 | | | 1.775 | x |

Table S3: Pairwise comparisons of wt-HepI and mt-HepI structures at various time points, with either the N or C terminal domains being aligned

| Time span | Starting | End | RMS | Domain |
|---|---|---|---|---|
| 700 | 1 | 160 | 2.437 | N |
| 800 | 1 | 160 | 2.926 | N |
| 900 | 1 | 160 | 2.663 | N |
| 1000 | 1 | 160 | 2.439 | N |
| 700 | 180 | 322 | 3.576 | C |
| 800 | 180 | 322 | 4.1 | C |
| 900 | 180 | 322 | 4.6 | C |
| 1000 | 180 | 322 | 4.399 | C |

SI Figure legends:

Figure S1: Phylogenetic tree of reference species created from marker genes.

Figure S2: RMSD of wt-HepI (black line) and mt-HepI (red line) till the end of the simulation.

Figure S3: RMSF of Cα for wt-HepI (black line) and mt-HepI (red line) from the converged trajectory.

Figure S4: RMSD for the pairwise comparisons by superimposition of structures at the various time points from converged trajectory for wt-HepI (upper panel) and mt-HepI (lower panel).

Figure S5: Hydrogen bonding network analysis of wild-type (wt-HepI) and mutant (mt-HepI) structures at selected time points from the converged trajectory. Each HepI position is represented on the circle's perimeter, with lines indicating hydrogen bonds: black (wt-HepI), red (mt-HepI), and yellow (both).

Figure S6: Secondary structural analysis followed by histogram distribution of wt-HepI (a-b) and mt-HepI (c-d) from the converged trajectory.

Figure S7: The projection of the first two eigenvectors (ev1 vs ev2) was conducted for (a) wt-HepI and (b) mt-HepI proteins.

Figure S8: Comparison of the motions in the two DCC matrices reveals that few regions retain their correlated motion patterns (red or blue), while many regions exhibit opposite correlation patterns between the two simulations (magenta).

Figure S1

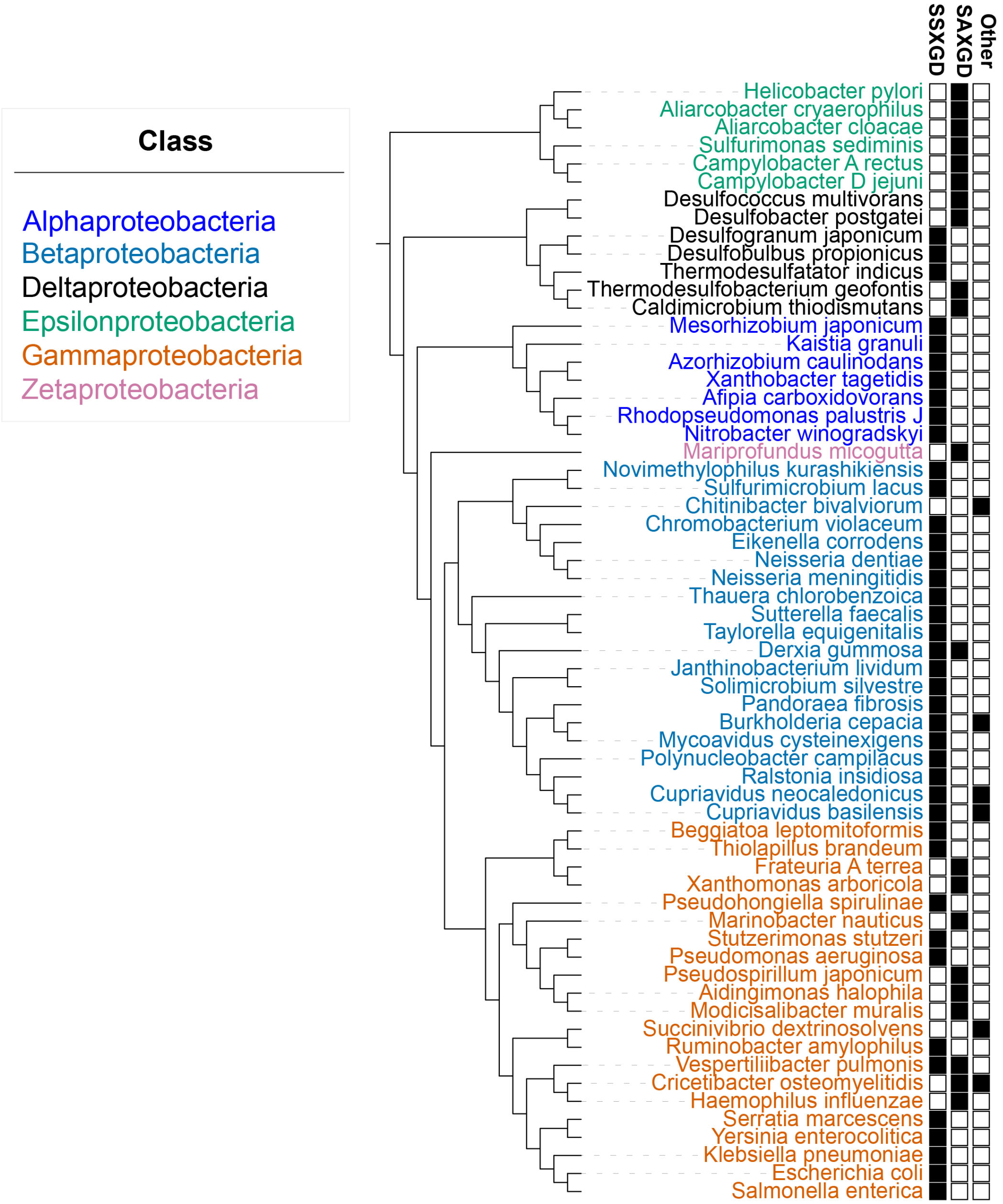
Class
Alphaproteobacteria
Betaproteobacteria
Deltaproteobacteria
Epsilonproteobacteria
Gammaproteobacteria
Zetaproteobacteria
SSXGD
SAXGD
Other
Helicobacter pylori
Aliarcobacter cryaerophilus
Aliarcobacter cloacae
Sulfurimonas sediminis
Campylobacter A rectus
Campylobacter D jejuni
Desulfococcus multivorans
Desulfobacter postgatei
Desulfogranum japonicum
Desulfobulbus propionicus
Thermodesulfatator indicus
Thermodesulfobacterium geofontis
Caldimicrobium thiodismutans
Mesorhizobium japonicum
Kaistia granuli
Azorhizobium caulinodans
Xanthobacter tagetidis
Afipia carboxidovorans
Rhodopseudomonas palustris J
Nitrobacter winogradskyi
Mariprofundus micogutta
Novimethylophilus kurashikiensis
Sulfurimicrobium lacus
Chitinibacter bivalviorum
Chromobacterium violaceum
Eikenella corrodens
Neisseria dentiae
Neisseria meningitidis
Thauera chlorobenzoica
Sutterella faecalis
Taylorella equigenitalis
Derxia gummosa
Janthinobacterium lividum
Solimicrobium silvestre
Pandoraea fibrosis
Burkholderia cepacia
Mycoavidus cysteinexigens
Polynucleobacter campilacus
Ralstonia insidiosa
Cupriavidus neocaledonicus
Cupriavidus basilensis
Beggiatoa leptomitoformis
Thiolapillus brandeum
Frateuria A terrea
Xanthomonas arboricola
Pseudohongiella spirulinae
Marinobacter nauticus
Stutzerimonas stutzeri
Pseudomonas aeruginosa
Pseudospirillum japonicum
Aidingimonas halophila
Modicisalibacter muralis
Succinivibrio dextrinosolvens
Ruminobacter amylophilus
Vespertiliibacter pulmonis
Cricetibacter osteomyelitidis
Haemophilus influenzae
Serratia marcescens
Yersinia enterocolitica
Klebsiella pneumoniae
Escherichia coli
Salmonella enterica

Figure S2

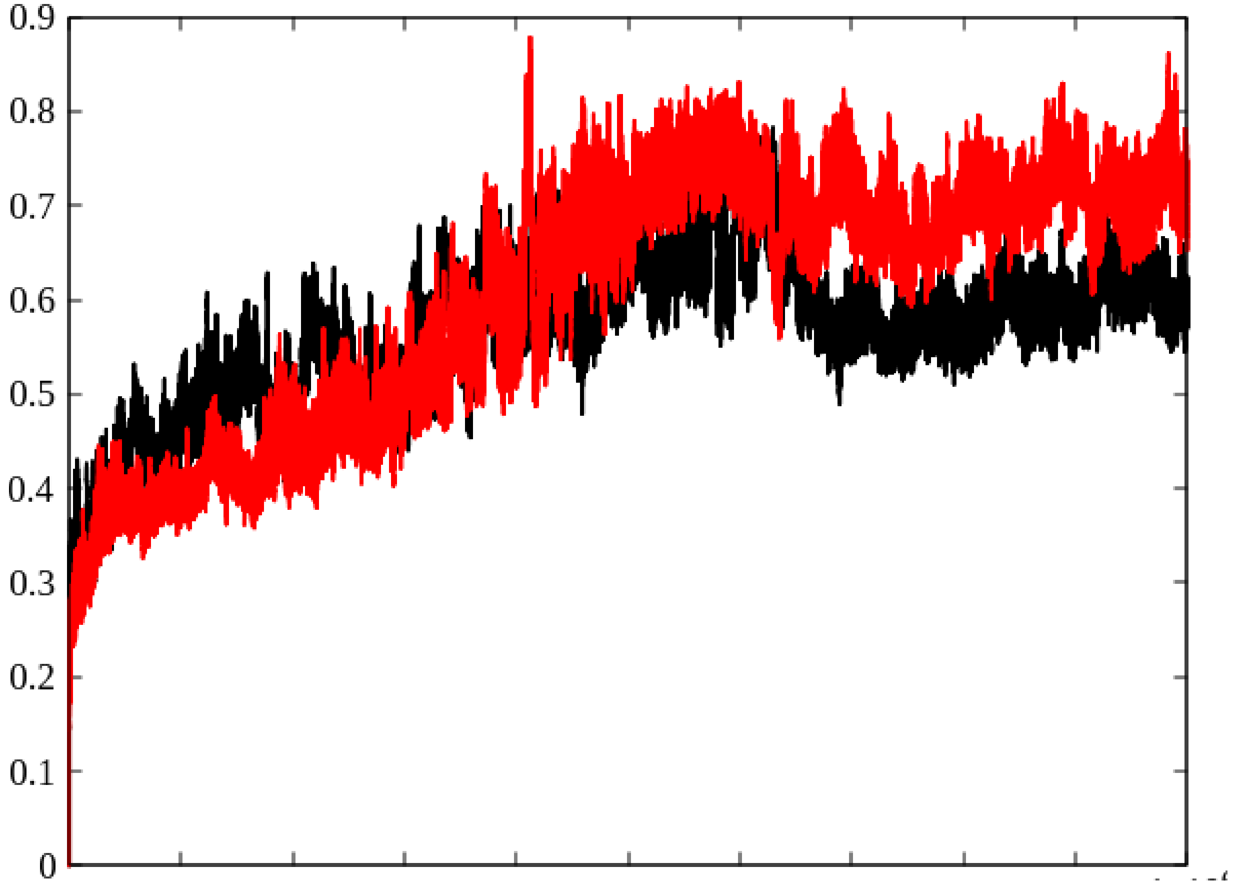

Figure S3

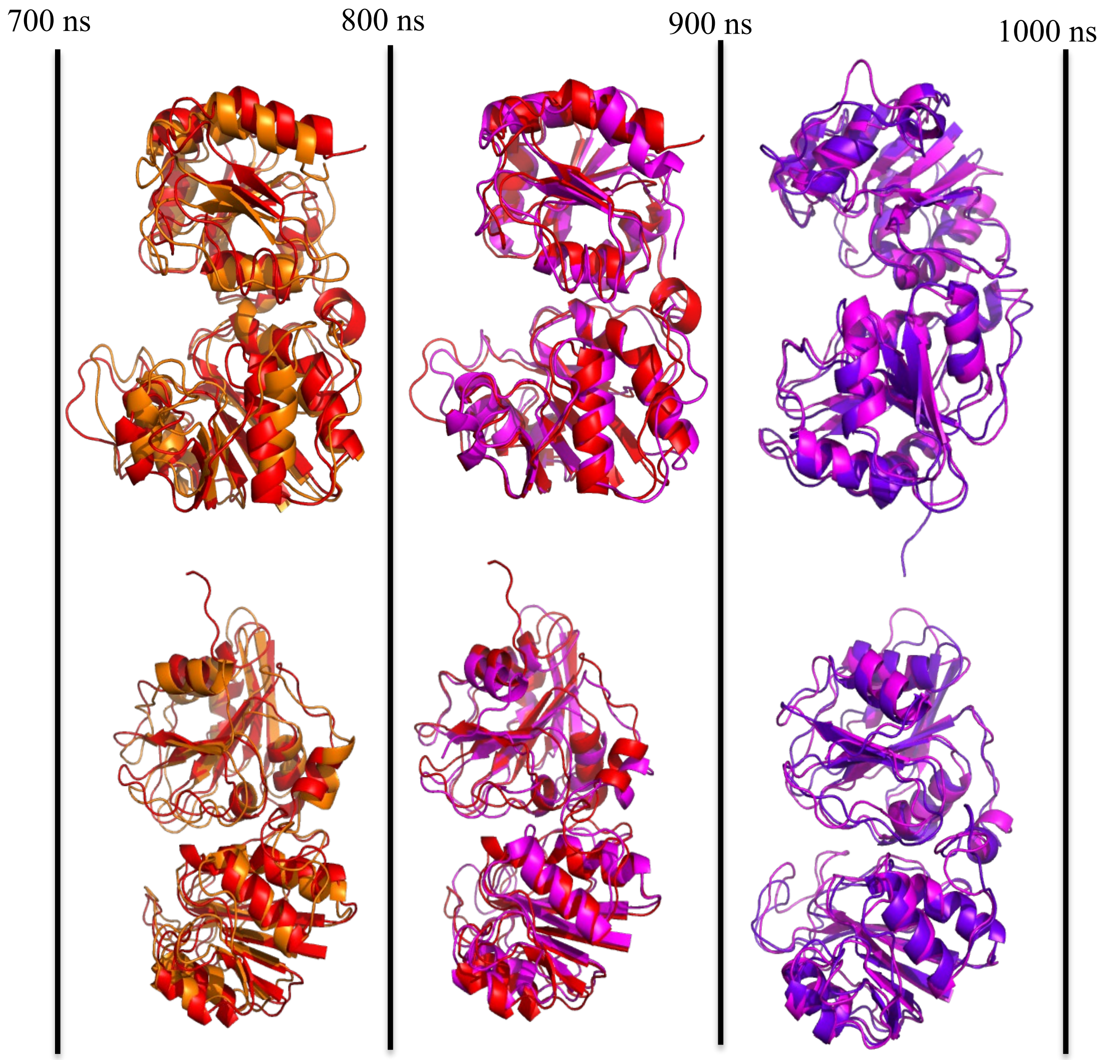

Figure S4

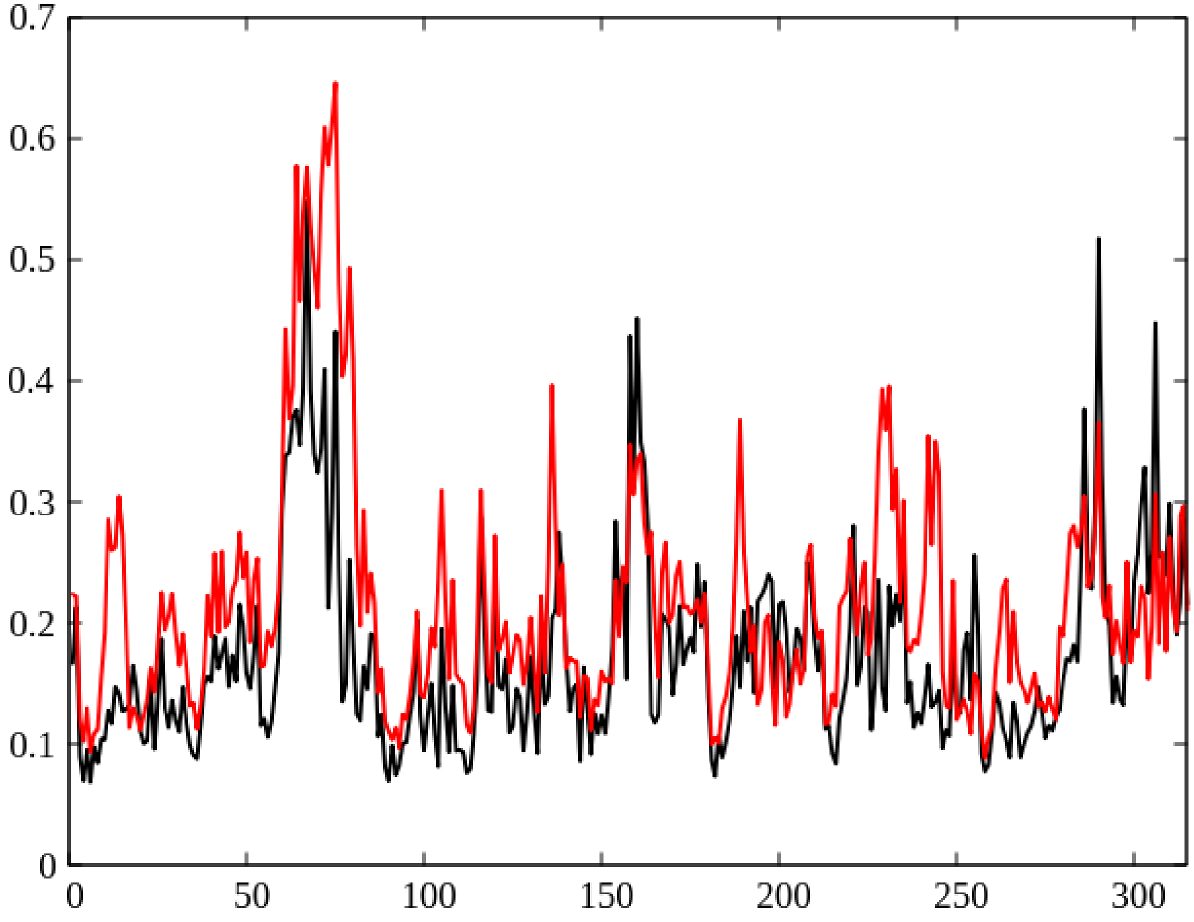

Figure S5

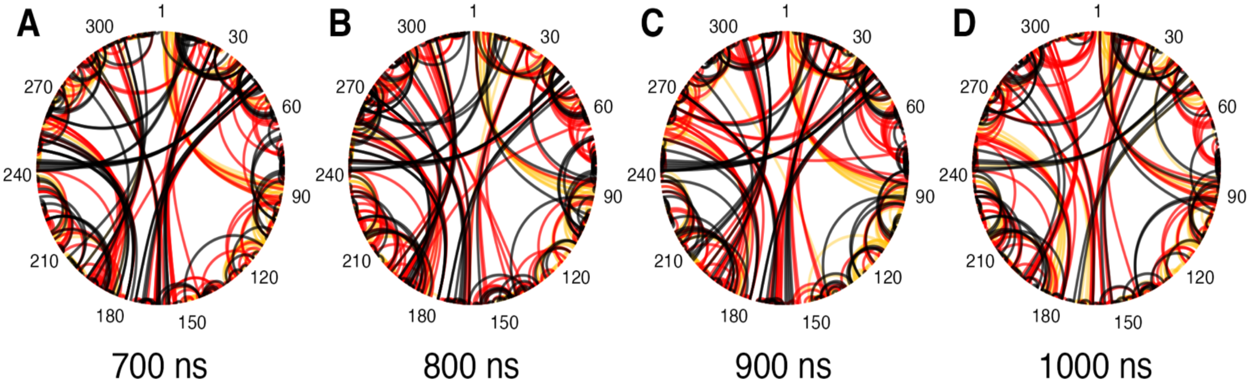
A
B
C
D
1
30
60
90
120
150
180
210
240
270
300
700 ns
800 ns
900 ns
1000 ns

Figure S6

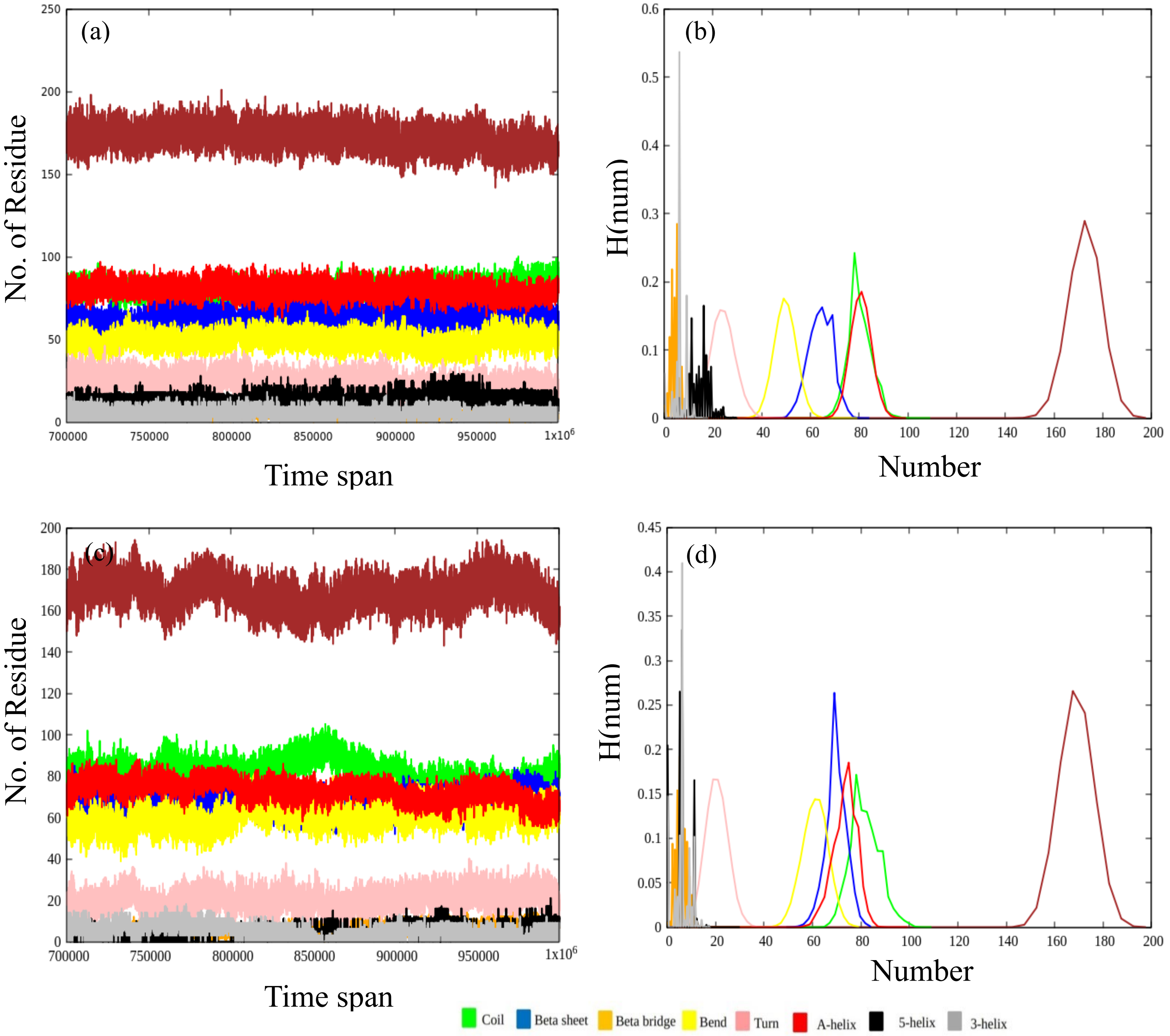

(a)
No. of Residue
Time span
(b)
H(num)
Number
(c)
No. of Residue
Time span
(d)
H(num)
Number
Coil
Beta sheet
Beta bridge
Bend
Turn
A-helix
5-helix
3-helix

Figure S7

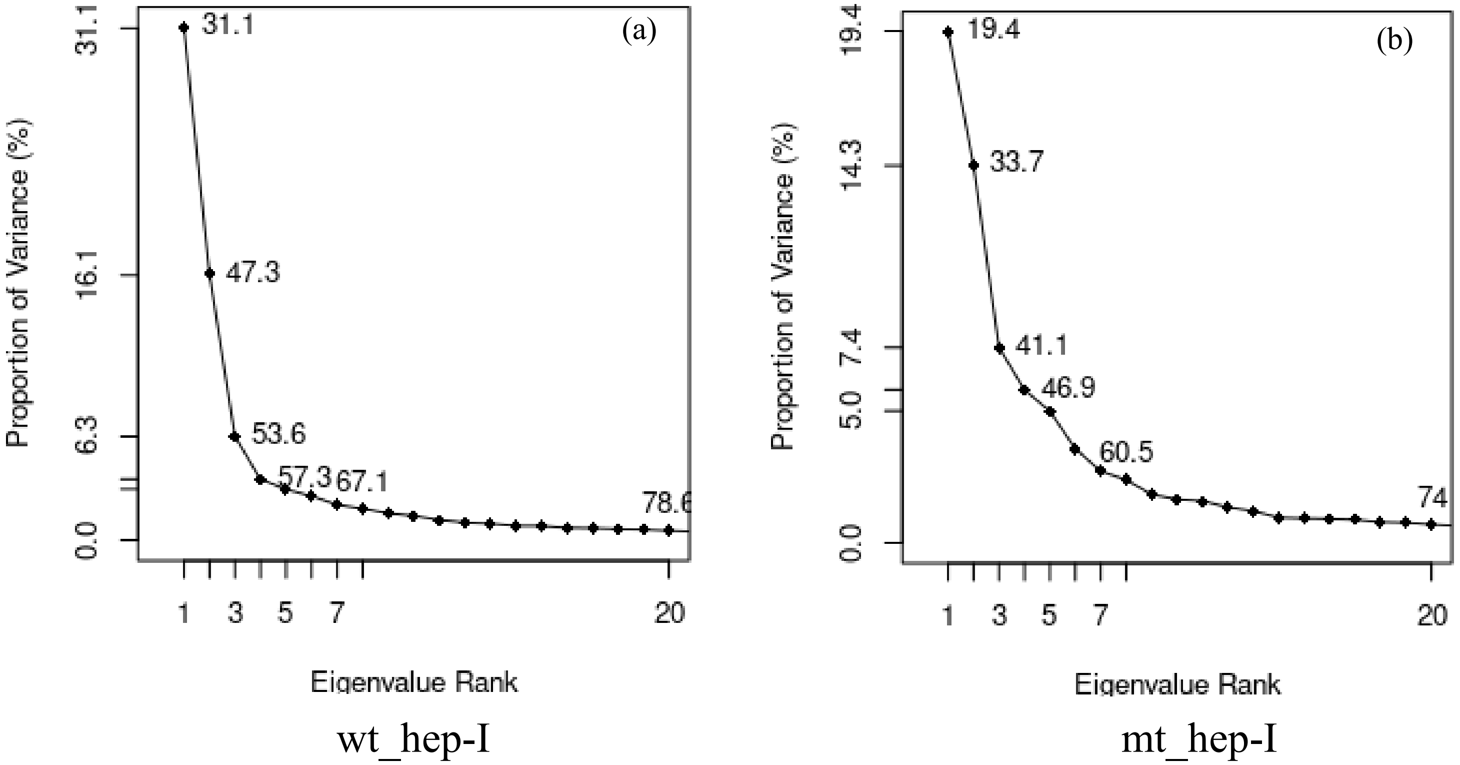

(a)
Proportion of Variance (%)
31.1
16.1
6.3
0.0
31.1
47.3
53.6
57.3
67.1
78.6
1
3
5
7
20
Eigenvalue Rank
wt_hep-I
(b)
Proportion of Variance (%)
19.4
14.3
7.4
5.0
0.0
19.4
33.7
41.1
46.9
60.5
74
1
3
5
7
20
Eigenvalue Rank
mt_hep-I

Figure S8

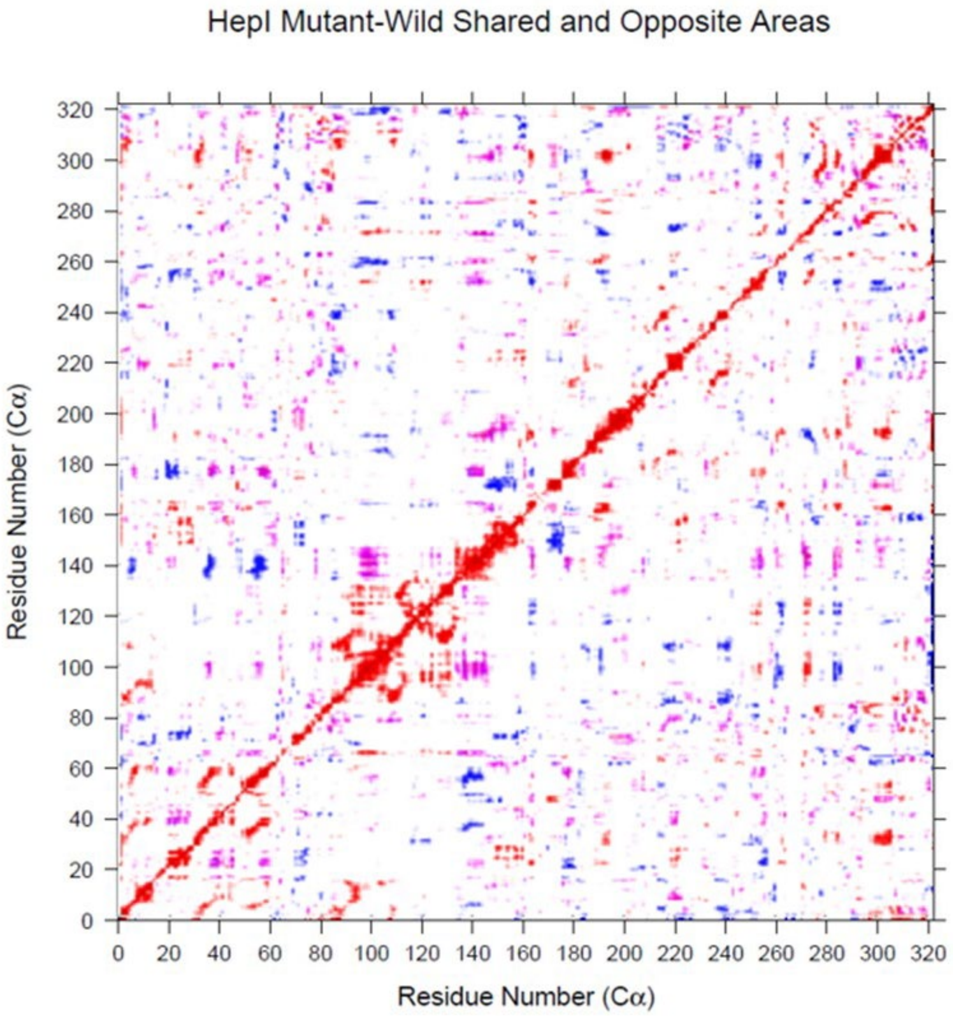
HepI Mutant-Wild Shared and Opposite Areas
Residue Number (Cα)
Residue Number (Cα)
0
20
40
60
80
100
120
140
160
180
200
220
240
260
280
300
320
